# Cell responses to simulated microgravity and hydrodynamic stress can be distinguished by using comparative transcriptomic analysis

**DOI:** 10.1101/2021.10.20.465119

**Authors:** Nikolai V. Kouznetsov

**Affiliations:** Karolinska Institutet, Solnavägen 9, SE-171 77 Stockholm, Sweden

**Keywords:** microgravity, hydrodynamic stress, T cell, gene expression, transcriptome, actin cytoskeleton networks

## Abstract

The human immune system is compromised in microgravity (MG) conditions during an orbital flight and upon return to Earth. T cells are critical for the immune response and execute their functions via actin mediated immune cell-cell interactions that could be disturbed by MG conditions. Here, we have applied two rotational platforms to simulate MG conditions: fast rotating clinostat (CL) and random positioning machine (RPM) followed by global T cell transcriptome analysis using RNA sequencing. We demonstrate that the T cell transcriptome profile in response to simulated MG treatment was clearly distinguishable from the T cell transcriptome response to hydrodynamic stress (HS) induced by shear forces upon cell movement in cultural medium. Gene expression profiling of genes related to or involved in actin cytoskeleton networks using RT-qPCR confirmed two sets of differentially regulated genes in the T cell response to MG or to HS. Several key genes potentially involved in T cell gravisensing (Fam163b, Dnph1, Trim34, Upk-1b) were identified. A number of candidate biomarker genes of the response to MG (VAV1, VAV2, VAV3, and NFATC2) and of the response to HS (ITGAL, ITGB1, ITGB2, RAC1 and RAC2) could be used to distinguish between these processes on the gene transcription level. Together, MG induces changes in the overall transcriptome of T cells leading to specific shifts in expression of cytoskeletal network genes.

## Introduction

The nature of gravitational sensor structures or mechanisms that respond to gravity change in T lymphocytes (T cells) is still to be determined. T cells need to rearrange their actin cytoskeleton for proper activation of the immune response [1].

The actin cytoskeleton proteins are putative candidates to participate in gravitational sensing [2, 3]. Microgravity induces the repression of T cell mediated immune response. Particularly, the T cell response towards mitogens in astronauts is reduced after space flight [2]. Human T cells stimulated with the mitogen concanavalin A reveal considerably reduced *in vitro* activation and cell proliferation in microgravity [2] and on a sounding rocket flight [4]. Stimulated T cells, pre-exposed to microgravity, show decreased expression of activation receptors including CD25, CD69 and CD71 and reduced inflammatory cytokine secretion and cell proliferation when compared to T cells under normal gravity [5]. Moreover, human lymphocytes undergo apoptosis upon exposure to modelled low gravity [6]. The inhibition of lymphocyte proliferation in microgravity is due to alterations occurring within the first hours of exposure to microgravity [7, 8]. Microarray expression analysis reveal changes in cytoskeletal gene expression [9] and overall altered patterns of global gene expression in space flown human T cells when compared to T cells activated in 1g control for normal gravity during space flight [10, 11]. Impaired induction of early genes, regulated by transcription factors NF-kB, CREB, ELK, AP-1 and STAT, contributed to T cell inhibition in artificial low gravity [12]. Analysis of gene expression in T cells from four human donors on board the International Space station (ISS) identified 99 down-regulated genes in microgravity [10]. In the context of cytoskeletal regulation, it was especially interesting that a majority of the genes that were inhibited in T cells in microgravity contains DNA serum response elements (SRE) in the promoters that binds the serum response factor (SRF) [10] as a part of transcriptionally active SRF/MRTFA complex. The SRF/MRTFA transcription factor plays a key role in actin cytoskeleton organization by targeting the transcription of genes encoding cytoskeletal proteins and is involved in several critical processes including cell growth and motility, apoptosis and cancer progression [13-15].

To explore changes in gene regulation of T cells under simulated microgravity during longer time points, we have applied the well-established microgravity simulation platforms at the German Aerospace Center (DLR). We compared T cells for 8 and 24 hours using clinostat (CL) and the random positioning machine (RPM) as key experimental systems [16] to simulate microgravity for suspension cell cultures at ground-based wet laboratory conditions (Figure 1A; Figure S1). We demonstrate that simulated MG (SMG) affects T cell gene expression profiles, particularly, the expression of actin cytoskeleton networks genes. SMG conditions induce shifts in the T cell transcriptome landscape, that can be clearly distinguished from characteristic transcriptome profile caused by hydrodynamic stress (HS) treatment. Several regulatory key genes potentially acting as components of T cell gravitation sensory pathways and a number of candidate marker genes of the response to MG and of the response to HS were identified.

**Figure 1.**
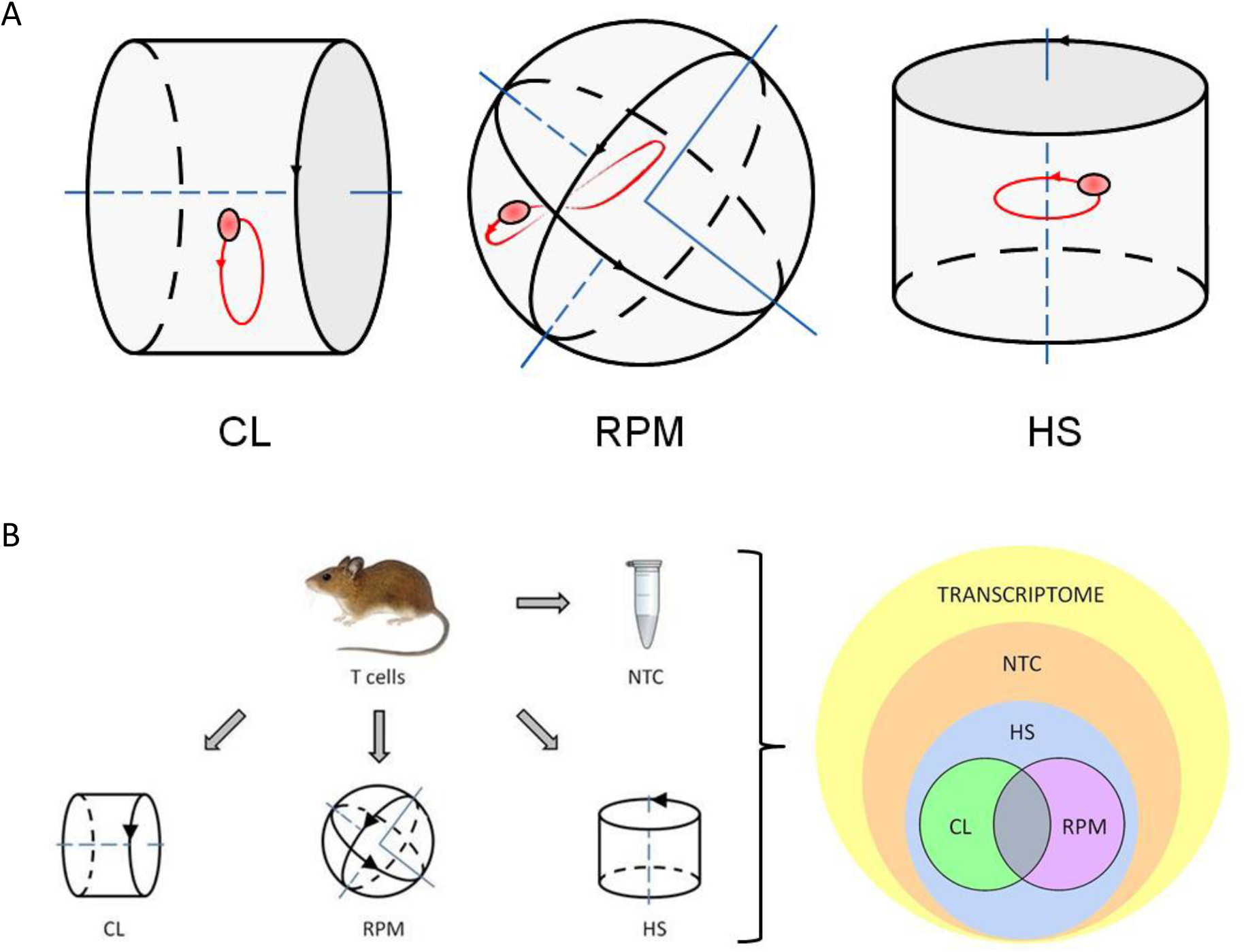
Experiment settings and principle of experiment. **A. The principle of simulated microgravity experiments**. Fast rotating clinostat (CL) and random positioning machine (RPM) used as simulated microgravity platforms. When rotating at sufficient speed, the suspension culture cells will be distributed homogeneously because the directional response to gravity is cancelled out. The cells rotate about themselves in simulated microgravity around one axis in Cl and move randomly around two axes in RPM. Orbital shaker used as a hydrodynamic cell stress inducing platform (HS). Direction of platform rotation is indicated by black arrows and cell suspension sample movement illustrated by exemplary single cell trajectory shown in red. **B. The overview chart of experiment settings and output of RNA-seq transcriptome data analysis**. Cell suspension of isolated primary mouse thymocytes (mouse T cells) was split into four parts with each part treated as indicated. All four parts were processed for RNA isolation and generation of RNA library followed by RNA sequencing. The output sequencing results showed that most of RNA transcripts of mouse T cell transcriptome were detected in SMG treated samples as well as in non-treated control (NTC) sample. Mouse T cell transcriptome response to microgravity in each of used experimental platforms - clinostat (CL) and random positioning machine (RPM) develops in parallel with and mainly on a transcriptional background of T cell response to hydrodynamic stress (HS). The set of transcripts in overlapping between CL and RPM panels were considered to represent gene candidates of the specific mouse T cell transcriptome response to simulated microgravity.

## Results

To setup the simulated microgravity (SMG) experiment, cell suspensions of isolated primary mouse thymocytes (mouse T cells) were subjected to hydrodynamic stress or exposed to SMG in a clinostat (CL) or random positioning machine (RPM) After exposure to these conditions the cells were processed for RNA isolation and generation of RNA library followed by RNA sequencing (RNA-seq). The differential gene expression analysis revealed arrays of genes differentially expressed in each of the experimental platforms - clinostat (CL), random positioning machine (RPM) and hydrodynamic stress (HS) inducing system. The set of transcripts shared between CL and RPM panels were considered to represent gene candidates of the specific mouse T cell transcriptome response to simulated microgravity (Figures 1B, 3, 4).

Overall transcript expression levels showed a similar distribution range in different RNA-seq samples with high coincidence between biological replicates (Figure 2A). In hierarchical clustering of differentially expressed genes (DEG), similar expression patterns were observed in samples after 8 hours in SMG conditions and it was possible to discern distinct pattern of DEGs between SMG and HS treated samples (Figure 2B).

**Figure 2.**
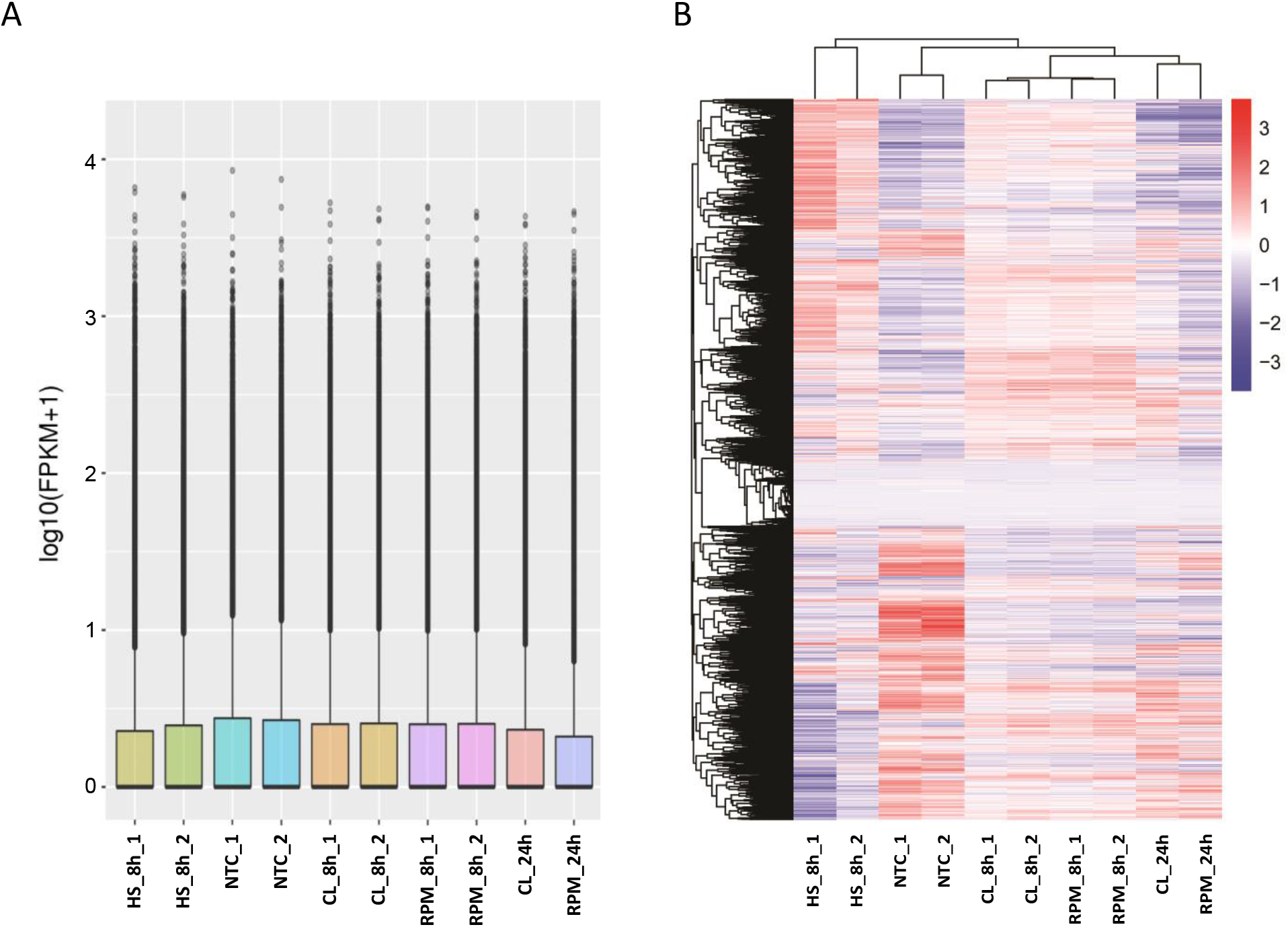
The overview of global transcriptome expression in RNA-seq samples. **A. FPKM distribution in RNA-seq samples** Boxplot demonstrates the FPKM distribution in RNA-seq samples, the x-axis shows the sample names and the y-axis shows the log_10_(FPKM+1) value. **B. Heatmap of clustering of differential gene expression in RNA-seq samples** The hierarchical cluster analysis was used to observe gene expression patterns under SMG or HS experimental conditions versus NTC. The colour scale from red to blue represents the log_10_(FPKM+1) value from high to low gene expression levels.

The number of terms in the Gene Ontology (GO) function analysis of corresponding genes and GO enrichment analysis was considerably lower when comparing SMG to HS treatment in contrast to the comparison of SMG to non-treated control (NTC) samples. This was true for all three categories of GO enrichment analysis: cellular component (CC), molecular function (MF) and biological processes (BP) (Figure S2). For transcription regulation, counts of molecular function term, “transcription factor binding, GO:0008134”, were 271 in CL vs NTC; 226 in RPM vs NTC and 35 in CL vs HS; 33 in RPM vs HS, respectively. Counts of the molecular function term, “DNA-binding transcription factor activity, GO:0003700”, were 279 in CL vs NTC; 247 in RPM vs NTC and 26 in CL vs HS; 30 in RPM vs HS, respectively. For organization of cytoskeleton, counts of the cellular component term, “cytoskeleton, GO:0005856 “, were 904 in CL vs NTC; 782 in RPM vs NTC and 132 in CL vs HS; 161 in RPM vs HS, respectively. Counts of the biological process term, “cytoskeleton organization, GO:0007010”, were 492 in CL vs NTC; 429 in RPM vs NTC and 74 in CL vs HS; 95 in RPM vs HS, respectively. Counts of molecular function term, “cytoskeletal protein binding, GO:0008092”, were 319 in CL vs NTC; 280 in RPM vs NTC and 49 in CL vs HS; 68 in RPM vs HS, respectively (Figure S2; Supplementary file 1). The top five common terms for RPM vs HS and CL vs HS at 8 hours of simulated microgravity are shown in Table 1 for three categories of GO enrichment analysis: cellular component (CC), molecular function (MF) and biological processes (BP).

**Table 1:** Top five common Gene Ontology (GO) terms at 8 hours of simulated microgravity in RPM and CL versus hydrodynamic stress (HS) treatment for three categories of GO enrichment analysis: cellular component (CC), molecular function (MF) and biological processes (BP).

| GO category | GO term ID | Description | Count, RPM vs HS, 8h | Count, CL vs HS, 8h | p-value, RPM vs HS, 8h | p-value, CL vs HS, 8h |
| --- | --- | --- | --- | --- | --- | --- |
| CC | 0043226 | organelle | 631 | 601 | 1.0982e-36 | 4.1886e-40 |
| CC | 0005737 | cytoplasm | 571 | 542 | 3.4794e-34 | 2.6777e-36 |
| CC | 0005634 | nucleus | 389 | 365 | 1.5571e-24 | 3.0424e-24 |
| CC | 0032991 | protein-containing complex | 355 | 363 | 1.001e-33 | 6.768e-44 |
| CC | 0005694 | chromosome | 120 | 97 | 7.3686e-27 | 4.2603e-17 |
| MF | 0019899 | enzyme binding | 136 | 120 | 2.0757e-10 | 0019899 |
| MF | 0005198 | structural molecule activity | 77 | 79 | 5.9878e-16 | 2.0991e-17 |
| MF | 0003723 | RNA binding | 72 | 87 | 4.4686e-06 | 0003723 |
| MF | 0008092 | cytoskeletal protein binding | 68 | 49 | 9.5479e-09 | 0008092 |
| MF | 0003735 | structural constituent of ribosome | 55 | 61 | 6.5833e-27 | 8.7689e-31 |
| BP | 0034641 | nitrogen compound metabolic process | 362 | 346 | 1.1454e-16 | 2.0049e-18 |
| BP | 0009058 | biosynthetic process | 306 | 288 | 1.6028e-10 | 6.6327e-11 |
| BP | 0007049 | cell cycle | 178 | 136 | 2.4143e-45 | 2.2782e-26 |
| BP | 0006950 | response to stress | 174 | 152 | 0.00015733 | 0.0079929 |
| BP | 0022607 | cellular component assembly | 136 | 133 | 3.6597e-06 | 1.3117e-07 |

In the co-expression analysis, the number of significantly co-expressed genes detected in common between the two biological replicates in the CL and RPM experiments at 8 hours of simulated MG were quite similar: 231 and 234 genes, respectively (Figure 3). To identify genes that were co-expressed in both CL and RPM platforms at 8 hours of simulated MG, the lists of 231 genes and 234 genes were compared and 113 common genes were identified (Supplementary file 2). Out of these 113 genes: 65 genes were known to be expressed in thymocytes (18 high or moderately expressed genes and 47 low expressed genes) and the remaining 48 genes were represented by pseudogenes, ncRNAs, and unknown sequences. Above 65 genes expressed in thymocytes were estimated for the significance of the transcript expression at 8 hours in simulated MG versus HS (Supplementary file 3). This estimation resulted in the list of seven genes found to be significantly differentially expressed, of which five were protein-coding genes expressed in murine thymocytes (Table 2).

**Table 2:** The significantly differentially expressed genes in CL or RPM or both platforms at 8 hours of simulated microgravity versus hydrodynamic stress treatment.

| Gene ID | Gene symbol | Gene name | T cell gene expression | Log2 Fold Change CL vs HS, 8h | Log2 Fold Change RPM vs HS, 8h |
| --- | --- | --- | --- | --- | --- |
| 109349 | Fam163b | family with sequence similarity 163, member B | + | -1,06 | -1,24 |
| 381101 | Dnph1 | 2'-deoxynucleoside 5'-phosphate N-hydrolase 1 | + | ns* | -1,44 |
| 434218 | Trim34b | tripartite motif-containing 34B | + | 2,83 | ns |
| 319581 | Xkr5 | X-linked Kx blood group related 5 | + | ns | -1,40 |
| 22268 | Upk1b | uroplakin-1b | + | -1,69 | ns |
| 6608222 | LOC6608222 | sodium/calcium exchanger protein | - | ns | -1,28 |
| - | Gm26737 | lncRNA, predicted gene, 26737 | - | ns | -1,55 |

\* ns - no significance

**Figure 3.**
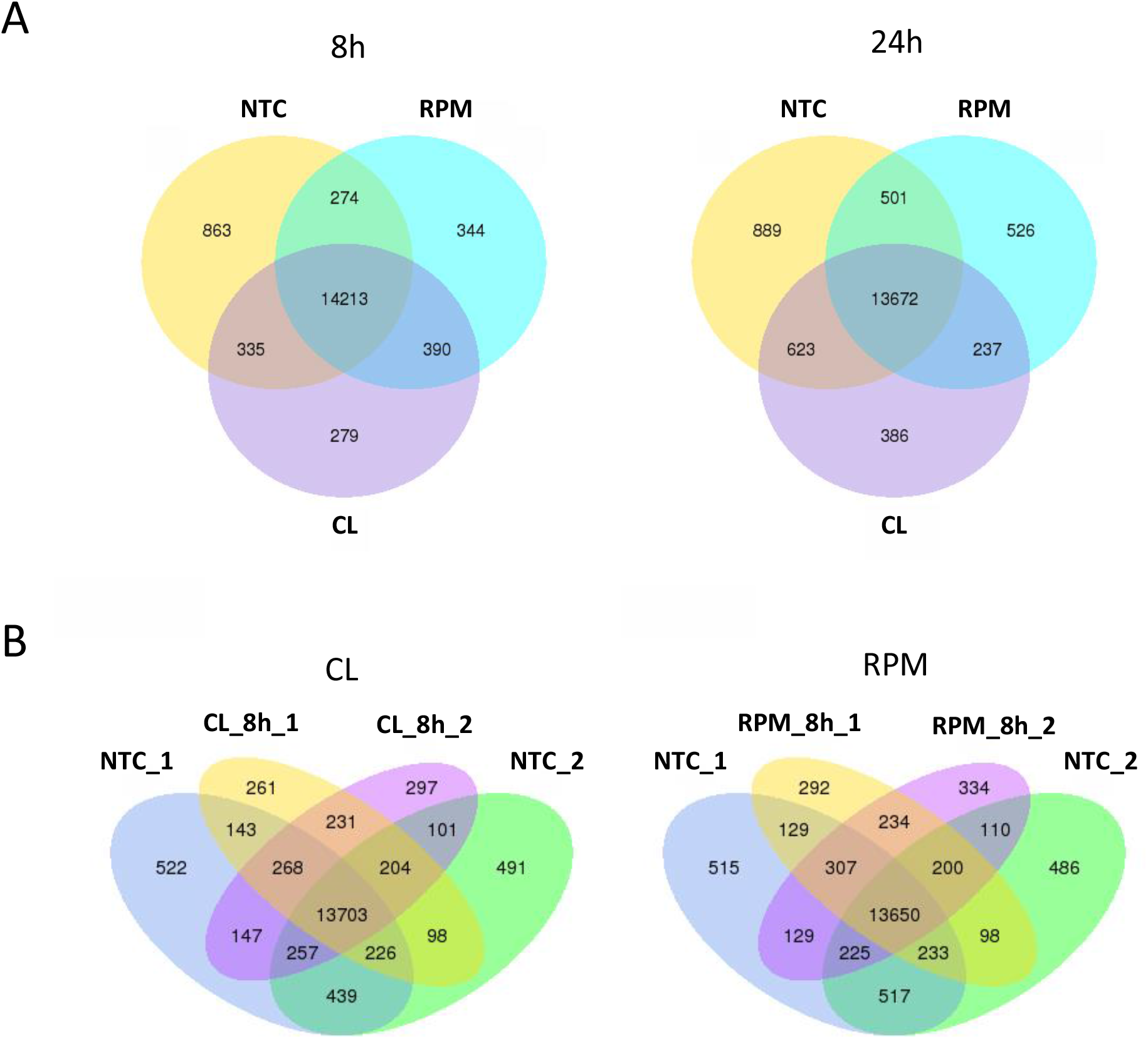
Venn diagrams indicating numbers of significantly expressed and co-expressed genes in simulated microgravity experiment. **A. Gene expression in simulated microgravity experiment at 8 hours and 24 hours** Numbers of expressed and co-expressed genes shown at 8 hours and 24 hours of treatment by simulated microgravity in clinostat (CL) and random positioning machine (RPM) versus non-treated control (NTC). **B. Gene expression in biological duplicates at 8 hours of simulated microgravity** Numbers of expressed and co-expressed genes indicated in biological duplicates at 8 hours in clinostat (CL) experiment (CL_8h_1, CL_8h_2) and random positioning machine (RPM) experiment (RPM_8h_1, RPM_8h_2) versus biological duplicates of non-treated control (NTC_1, NTC_2).

Fam163b with unknown function was the only significantly down-regulated gene in both CL and RPM samples vs HS condition. DEGs down-regulated at 8h in RPM vs HS included 2’-deoxynucleoside 5’-phosphate N-hydrolase 1 (Dnph1), also known as RCL; C6orf108; dJ330M21.3, a putative oncogene with a role in cellular proliferation and c-Myc-mediated transformation [17]. Interestingly, DNPH1 was shown to be down-regulated post-flight in the NASA twins study [18]. Another gene in this group was X-linked Kx blood group related 5 (Xkr5), also known as XRG5; 5430438H03Rik gene, involved in the engulfment of apoptotic cell corpses [19]. Two other significantly expressed DEGs: tripartite motif-containing 34 (Trim34) and uroplakin-1b (Upk-1b) were found to be up-regulated and down-regulated at 8h in CL vs HS, respectively. The human homolog TRIM34, was shown to facilitate the formation of multinucleated giant cells by enhancing cell fusion [20]. Upk-1b, Uroplakin-1b, also known as Tsp, is a member of the tetraspanin family. Most tetraspanins are cell-surface proteins that mediate signal transduction in the regulation of cell development, activation, growth, cell migration and motility [21, 22].

As was the case with the GO enrichment analysis, the number of genes in the DEG analysis was considerably lower when comparing MG treatment and HS control samples versus comparing MG treatment and non-treated controls (NTC). The number of upregulated/downregulated transcripts were 255 and 700 in CL versus HS; 284 and 771 in RPM versus HS; 5211 and 1644 in CL versus NTC; 4534 and 1704 in RPM versus NTC, respectively (Figure 4A-B).

**Figure 4.**
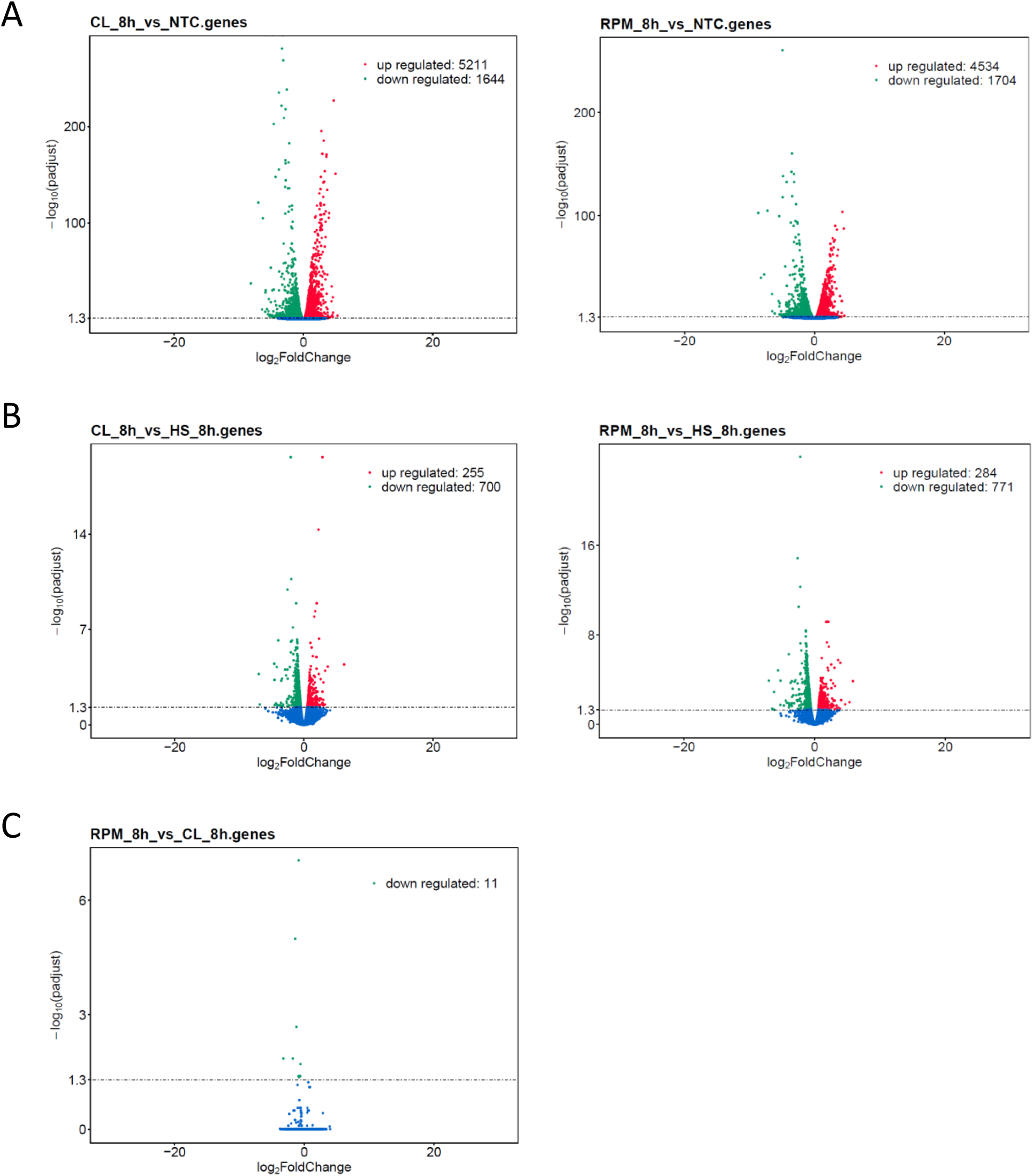
Volcano plots displaying overall distribution of differentially expressed genes in murine T cells at 8 hours of simulated microgravity. The x-axis shows the fold change in gene expression, and the y-axis designates the statistical significance of the differences between treatment group and control group with the threshold set as: padj < 0.05. Significantly up- and downregulated genes highlighted in red and green, respectively, and other genes shown in blue. Treatment group versus control group data illustrated for comparisons: **A. Treatment by simulated microgravity in clinostat (CL) and random positioning machine (RPM) versus non-treated control (NTC);** **B. Treatment by simulated microgravity in clinostat (CL) and random positioning machine (RPM) versus hydrodynamic stress (HS);** **C. Treatment by simulated microgravity in random positioning machine (RPM) versus clinostat (CL)**.

In addition, DEG analysis revealed 11 downregulated transcripts in RPM platform versus CL platform that represent a unique gene signature which discriminates SMG treatments in RPM and CL (Figure 4C, Table 3).

**Table 3:** The significantly down-regulated genes in CL platform versus RPM platform at 8 hours of simulated microgravity treatment.

| Gene ID | Gene symbol | Gene name | T cell gene expression | Log2 Fold Change RPM vs CL, 8h | Log2 Fold Change CL/RPM vs HS, 8h |
| --- | --- | --- | --- | --- | --- |
| 68195 | Rnaset2b | ribonuclease T2B | + | -0,87 | ns / ns* |
| 16149 | Cd74 | CD74 antigen | + | -0,69 | ns / -1.2 |
| 14086 | Fscn1 | fascin actin-bundling protein 1 | + | -1,21 | 0.82 / ns |
| 208084 | Pif1 | PIF1 5'-to-3' DNA helicase | + | -0,61 | ns / -0.83 |
| 108956 | Apol7c | apolipoprotein L 7c | + | -1,69 | 2.86 / 1.46 |
| 100042807 | Eif3j2 | eukaryotic translation initiation factor 3 s.J2 | + | -1,75 | ns / ns |
| 193740 | Hspa1a | heat shock protein 1A | + | -3,24 | ns / ns |
| 12316 | Aspm | abnormal spindle microtubule assembly | + | -0,57 | ns / ns |
| 17476 | Mpeg1 | macrophage expressed gene 1 | + | -0,90 | ns / ns |
| 14969 | H2-Eb1 | histocompatibility 2, class II antigen E beta | + | -0,80 | ns / -1.23 |
| 432447 | Gm9824 | predicted pseudogene 9824 | - | -0,81 | ns / -0.99 |

\* ns - no significance

Several of these genes are conserved in higher Vertebrata and expressed in mouse and human T cells and are of particular interest: Hspa1a, Cd74, Fscn1, Rnaset2b, Pif1. Ubiquitously expressed chaperone heat shock protein 1A (also known as Hspa1a, Hsp; Hsp7; Hsp70) stabilizes existing proteins against aggregation and mediates the folding of newly translated proteins [23]. The CD74 antigen (CLIP, DHLAG, etc.) also serves as an important chaperone in immune cells and plays a role in initiation of survival pathways and cell proliferation [24]. Fascin actin-bundling protein 1 (fascin-1, Fscn1, Fan1) is involved in actin cytoskeleton organization and biogenesis and acts within actin filament bundle assembly. The encoded protein plays a critical role in cell migration, motility, adhesion and cellular interactions [25, 26]. Ribonuclease T2 (Rnaset2b) regulates mitochondrion-associated cytosolic ribosomes [27]. PIF1 5’-to-3’ DNA helicase (Pif1) resolves G-quadruplexes and RNA-DNA hybrids at the ends of chromosomes and, also, prevents telomere elongation by inhibiting the actions of telomerase [28, 29]. These and a few other genes were found specifically induced in the CL system (Figure 4C, Table 3).

The gene expression of selected 27 genes related to or involved in the actin cytoskeleton networks was compared in cultured human T cells (Jurkat) under simulated MG in both CL and RPM using RT-qPCR. The differential gene expression profiles showed similar patterns for most of the considered genes in both CL and RPM simulators (Figure 5A). Gene expression changes were compared between cells that were subjected to simulated MG and HS. The 27 genes were grouped into those in which the expression was greater, less than and equal to the expression in simulated MG when compared to the expression in HS. From this comparison of differential gene expression, it was possible to identify genes in which the change in expression was specific to either simulated MG or to HS (Figure 5B, Supplementary Table S1). Particularly, the members of the VAV family of guanine nucleotide exchange factors (VAV1, VAV2, VAV3) were downregulated in simulated MG while their expression was constant in HS. The VAV proteins mediate changes in the actin cytoskeleton via control of the small Rho GTPases [30, 31]. The expression of the NFATC2 gene encoding a transcription factor important for the development and differentiation of T cells [32, 33] was suppressed in simulated MG while unchanged in HS (Figure 5B). Candidate marker genes of the T cell response to HS were identified. Among these were genes encoding integrins that play a key role in intracellular adhesion and cell-cell interactions and members of the Rac family of small GTPases mediating changes in the actin cytoskeleton, for example, in cell adhesion and lamellipodium formation [34]. The expression of ITGAL, ITGB1, ITGB2, RAC1 and RAC2 was reduced in the cells under HS but not in cells in simulated MG. Therefore, differential gene expression changes specific to both the response to simulated MG and HS were identified. These genes could potentially be used as markers for the response of T cells to MG and to HS (Supplementary Figure S3).

**Figure 5.**
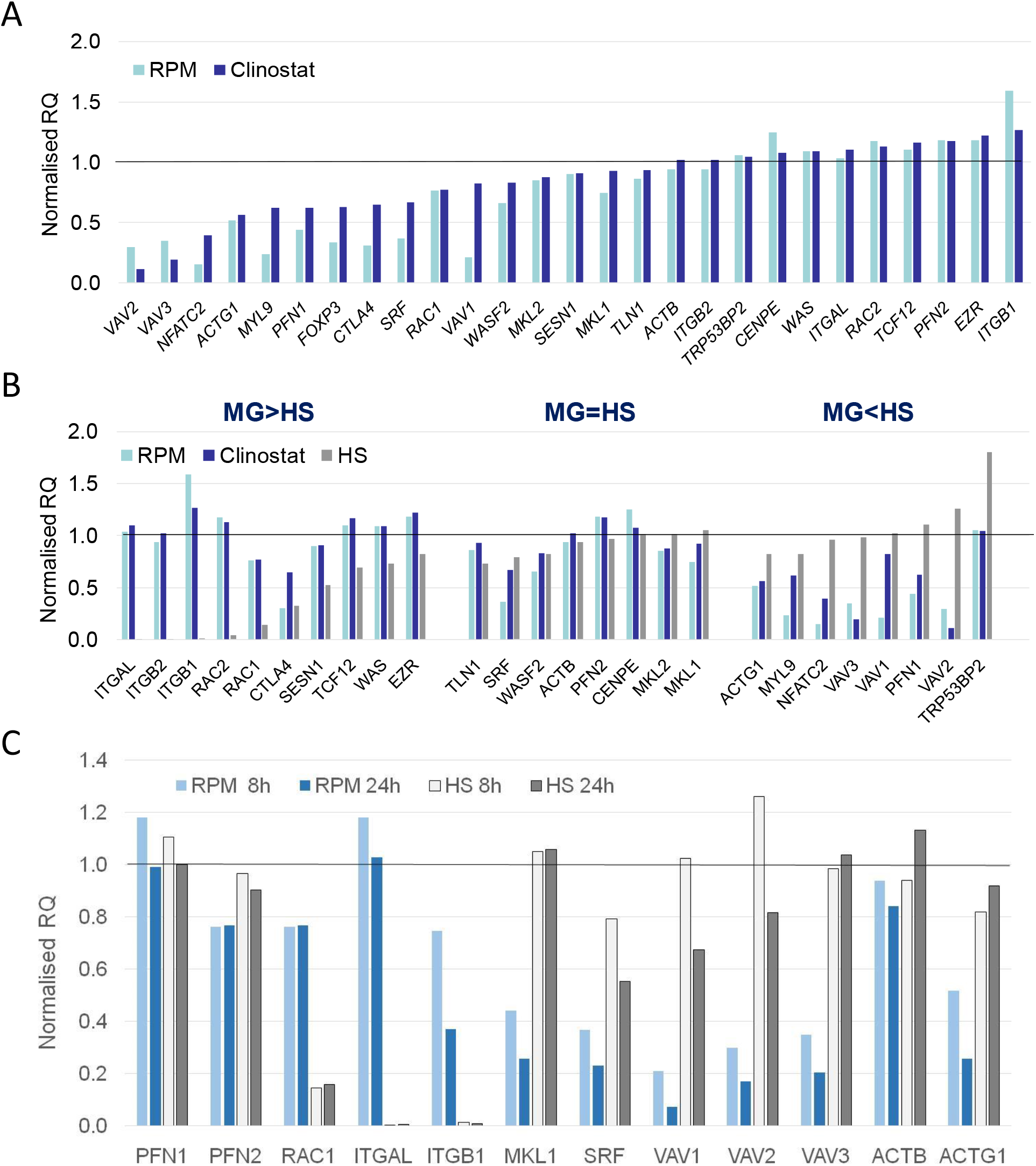
Differentially expressed genes in human T cells in simulated microgravity experiment measured by RT-qPCR. **A. Actin cytoskeleton network differential gene expression response to simulated microgravity** Differential expression of 27 genes measured at 8 hours of simulated microgravity in random positioning machine (RPM) or clinostat and normalised by values of non-treated control (RQ value equals 1.0). **B. Candidate marker genes discriminating T cell responses to simulated microgravity and to hydrodynamic stress** Differential expression measured at 8 hours of treatment in random positioning machine (RPM), clinostat or hydrodynamic cell stress inducing platform (HS). Genes grouped according to T cell responses in MG and HS for gene expression higher in simulated microgravity (MG>HS) or in hydrodynamic stress treatment (MG<HS) or in a similar range (MG=HS); **C. Dynamic expression of actin network selected key genes in simulated microgravity versus hydrodynamic stress response** Differential expression measured at 8 hours and 24 hours of treatment in random positioning machine (RPM) or hydrodynamic stress inducing platform (HS) and normalised by non-treated control values (RQ value equals 1.0).

Gene expression changes in Jurkat cells were also analysed at both 8 and 24 hours. This provided a dynamic view of gene expression and the trends in expression over time could be identified. The expression level of PFN1 and RAC1 was relatively constant in CL and RPM at both time points. However, RAC1 was further down-regulated in response to HS when compared to simulated MG. PFN1 encodes profilin 1, a regulator of actin filament formation [35]. The integrins ITGAL and ITGB1 showed the same trend in gene expression and were downregulated in response to HS. Under simulated MG, their expression decreased over time from 8 hour to 24 hour. MKL1/MRTFA expression was constant in response to HS but decreased in simulated MG and was further reduced over time from 8 hour to 24 hour. MKL1/MRTFA encodes an actin sensor that upon decreased G-actin enters into the nucleus and interacts with the serum response factor (SRF) to drive transcription of actin network genes [13, 14]. VAV1, VAV2 and VAV3 all followed the same trend in gene expression. There was a greater reduction in expression in response to simulated MG conditions while a decrease in expression was seen at 24 hours in both simulated MG and HS. All of these genes are involved in or closely related to regulation of the actin cytoskeleton. The expression of actin β and actin *γ*1 was analysed and compared at both 8 and 24 hours. This revealed an opposing trend in response to HS and simulated MG at the level of actin expression. Cells exposed to simulated MG, had downregulated ACTB and ACTG1 and the trend in time was a further reduction in expression. In contrast, there was an increase in the expression of ACTB and ACTG1 relative to the NTC which further increased between 8 and 24 hours.

Together, simulated MG and HS both have an influence on the actin cytoskeleton network of T cells at the level of gene expression. However, the response is in opposing directions with an increase in actin expression in cells under HS while actin expression levels decreased in simulated MG.

## Discussion

T cells constitute an important part of immune system that is affected by microgravity conditions and become compromised during or following the space flights in a large cohort of astronauts [7, 9, 11, 36].

Although not replacing the possibilities of expensive and rare experiments on the effects of weightlessness at Earth orbital facilities, the simulated microgravity (SMG) platforms proved to be valuable tools to study the nature of cellular mechanisms and gravisensing processes in ground based wet laboratory settings [16]. In this project, the T cell response to SMG conditions was studied at the level of gene expression in primary mouse T lymphocytes (thymocytes) and cultured human T lymphocytes (Jurkat) using two ground based applied systems: fast rotating clinostat (CL) and random positioning machine (RPM) (Supplementary Figure S1). Treatment was performed for 8-24 hours in sealed plastic tubes at 37oC in complete RPMI-1640 cell culture medium. In CL, the position of the sample is constantly changed around one axis with respect to the direction of Earth’s gravity vector and MG is simulated when the position of the sample changes faster than the sample can respond to gravity [37]. RPM uses the same underlying principle to simulate MG, however, rotation is performed around two axes in random mode (Figure 1A). [38].

In biological, *in vitro* experiments, non-treated control (NTC) traditionally is used as a best guess normaliser to analyse readout data. However, both rotational SMG platforms that were used employ forced cell movement in liquid. Therefore, an important consideration when using these MG simulators is that cell samples are rotating in a viscous cultural media exposing cells to shear forces which induces hydrodynamic stress (HS). HS was also found in microchannels of microfluidic and flow cytometry systems, in bioreactors where it has been shown to induce apoptosis and cause changes in the cytoskeleton of cells [39-42] and in the cells of tissues exposed to mechanical forces of physiological fluids [43-45].

Therefore, in order to distinguish between gene expression changes induced by MG from those caused by response to HS, T cells in MG simulators were compared to those subjected to HS alone in the same type of plastic tubes and culture medium at same temperature in a rotating orbital shaker for 8-24 hours (Figure 1B).

Inclusion of NTC and HS samples in the study as the parallel controls, showed that the use of HS for data normalisation decreased the transcriptome expression value noise considerably (Figure 4 A,B) and highlighted gene ontology (GO) terms that are specific for T cell transcriptome response to SMG (Supplementary Figure S2).

Since T cell cytoskeleton is involved in many functions of lymphocytes such as cell migration and cell-cell interactions, it is possible that the impact of microgravity on the cytoskeleton causes the immunosuppression experienced by the astronauts. Analysis of gene expression response at different timepoints demonstrates certain dynamics of T cell response to SMG. Moreover, a number of genes encoding constituents of actin cytoskeleton network genes changes expression in SMG with time that was confirmed by RT-qPCR (Figure 5C).

Finally, the response of T cell transcriptome to 8 hours of SMG treatment in clinostat (CL) and random positioning machine (RPM), appeared to have quite similar gene expression patterns (Figures 2B, 5A). However, a closer look at these responses and the comparison of differential expression at this timepointrevealed a panel of 10 genes known to be expressed in thymocytes. As these 10 genes were found to be upregulated in clinostat platform treatment versus RPM system (Figure 4C, Table 3), one may speculate, that their expression is inhibited, probably, due to higher shear stress that was previously observed in RPM when compared to clinostat conditions [46].

## Material and Methods

### Mouse primary T cells isolation

Wild-type mouse primary thymocytes (mouse T cells) were isolated from three C57BL/6 mouse thymi as sources for biological triplicates. Consistent measurements of one biological sample performed at least twice are considered as technical replicates. Thymi were homogenized through 100 μm strainers (Corning) and cells were resuspended in sterile 1x PBS. Mouse T cells were diluted to the final concentration 2 mln. cells / ml in sterile cultural medium RPMI-1640 (Gibco) containing 10% FBS and 0.1 M HEPES, pH 7.4. The resulting mouse T cell suspensions were used for all conditions and time points of experiment and in non-treated control.

### Cultured human T cells

An immortalised line of human T lymphocytes (human T cells) (Jurkat, Clone E6.1 TIB-152 (ATCC) was propagated from three frozen aliquots and subcultured cell stocks were used as sources for biological triplicates. Consistent measurements of one biological sample performed at least twice are considered as technical replicates. Human T cells were diluted to final concentration 2 mln. cells/ml in sterile cultural medium RPMI-1640 (Gibco) containing 10% FBS and 0.1 M HEPES, pH 7.4. The resulting human T cell suspensions were used for all conditions and time points of experiment and in non-treated control.

### Experimental platforms

The fast rotating clinostat (CL) and random positioning machine (RPM) were used in a comparative approach as ground-based facilities to simulate microgravity for suspension cell cultures of primary mouse T cells and cultured human T cells as described previously [16]. Applied regimes based on previous optimisation experiments were as follows: 60 rpm for CL, continuously rotating around one rotation axis perpendicular to the direction of the gravity vector (Supplementary Video 1), 12 rpm for RPM, consisting of 2 rotation axes with constant speed and uniform directional movement for RPM (Supplementary Video 2), and constantly 180 rpm for orbital hydrodynamic cell stress (HS) inducing platform continuously rotating around one rotation axis parallel to the direction of the gravity vector. HS inducing platform was loaded with cylindrical external containers with a radius of 14 mm and HS sample tubes were placed into containers (Supplementary Video 3).

In the clinostat, cell suspension samples were exposed in 1 ml polystyrene pipettes with a radius of 1 mm. Thus maximal residual acceleration of clinorotation did not exceed 0.04 g according to previously suggested formula and calculation [47], [48]. In the RPM and HS platforms and in stationary 1g non-treated control (NTC), cell samples were exposed in 5 ml polystyrene cylindrical tubes with a radius of 5 mm.

### RNA extraction

Total RNA was purified from sorted cells using RNeasy kit #74104 (QIAGEN). Each RNA sample was treated with DNA nuclease using RNAse-free DNAse kit # 79254 (QIAGEN) during RNA isolation procedure to avoid contamination by genomic DNA. The concentration of purified total RNA was measured by NanoDrop 2000 (Thermo Scientific). RNA integrity and purity was visually evaluated by 1% UltraPureAgarose, #16500500 (Invitrogen), 1xTAE gel electrophoresis analysis. Gel images were created with ImageQuant LAS 4000 LAS4000 Image system using ImageQuant LAS 4000 Control Software (GE Healthcare).

### Real-time qPCR-based gene expression profiling

Two μg of total RNA per sample was used for each cDNA synthesis reaction (with SuperScript™ II Reverse Transcriptase, #18064-014 (Invitrogen). RT-qPCR reactions were performed using SsoAdvanced™ Universal SYBR® Green Supermix kit, #1725272SP (Bio-Rad) with C1000/CFX96 RT-PCR System (Bio-Rad).

Gene specific primers were designed with Primer Bank software (Supplemetary file 4).

Gene expression profiling by real-time qPCR was performed as described previously [33] in three technical replicates for each cDNA template sample with Ct variation between triplicates less than cut-off +/-0.25. Gene expression data: the mean of the technical triplicate, ΔCt, RQ value and standard deviation between RQ of biological triplicates were calculated and analysed using Excel software.

### RNA-seq and bioinformatic analysis

RNA sequencing and data quality control was performed with Illumina HiSeq-PE150 Platform at Novogene, Hongkong (Suplementary Table 2).

Mapping to the reference genome was performed with HISAT2, v.2.0.5. Reads were aligned to the reference genome of the house mouse (Mus musculus) assembly December 2011 (GRCm38/mm10) (Suplementary Table 3).

Standard bioinformatics analysis of RNA-seq data was performed by Novogene (Hongkong) including primary data filtering to remove adaptor sequences, contamination, and low-quality reads from raw reads, reads alignment and genome-wide distribution of RNA-seq reads. Gene ontology (GO) function analysis of corresponding genes and GO enrichment analysis [49].

RNA-seq differential gene expression analysis was performed with DESeq2 in R package. In gene expression analysis, H-cluster, K-means and SOM were used to cluster the log_2_ (ratios) (Suplementary Table 4) [50-52].

Functional classification of refined data for peak-related genes was performed using NCBI Gene resource (Bethesda MD, 20894 USA) and gene list were made with Excel software.

## Acknowledgments

This work was supported by research grant from the Swedish National Space Agency in which N.V.K was a co-recipient. I thank colleagues at the Institute of Aerospace Medicine at German Aerospace Center in Cologne, Germany and Karolinska Institutet in Stockholm, Sweden who will be added as co-authors of this article in the final version of manuscript.

## Supplementary information

**Supplementary figure S1:**
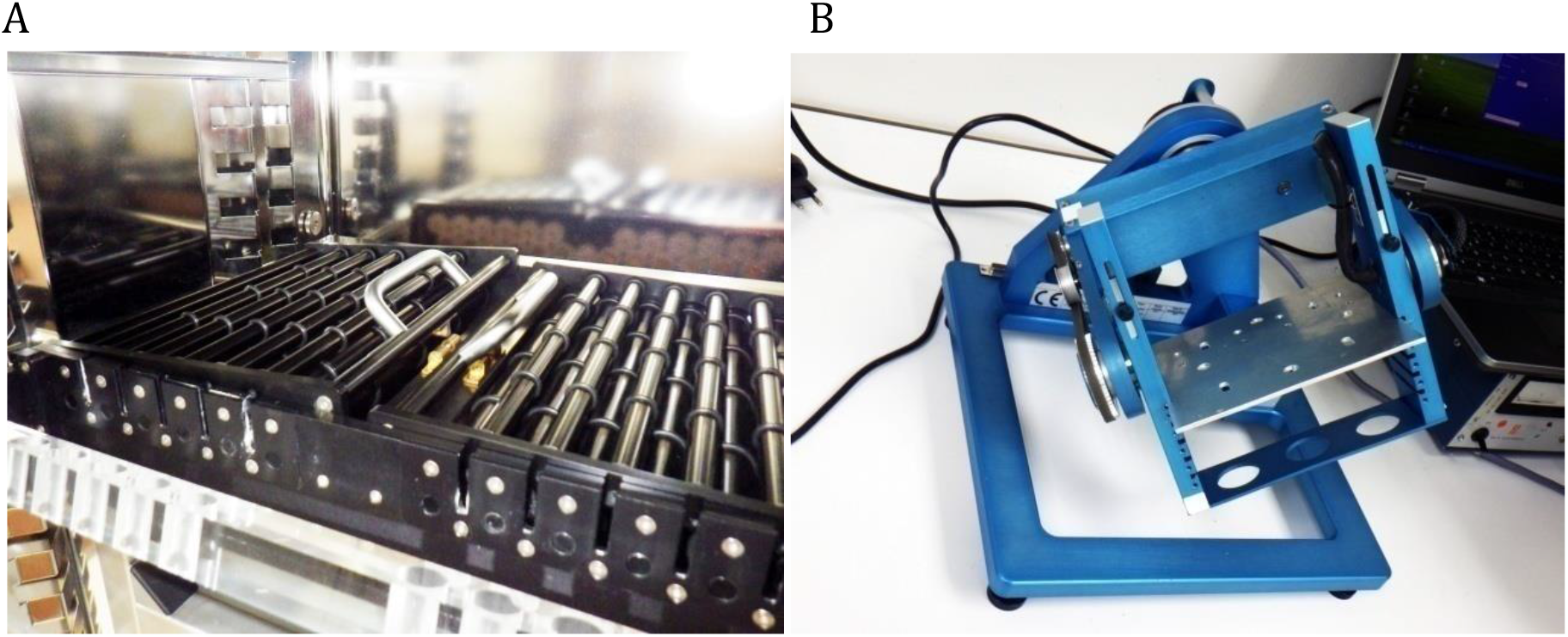
Simulated microgravity platforms (DLR) Microgravity simulation platforms (DLR) used as key experimental systems to simulate microgravity for suspension cell cultures: (A) Clinostat, (B) Random positioning machine.

**Supplementary figure S2:**
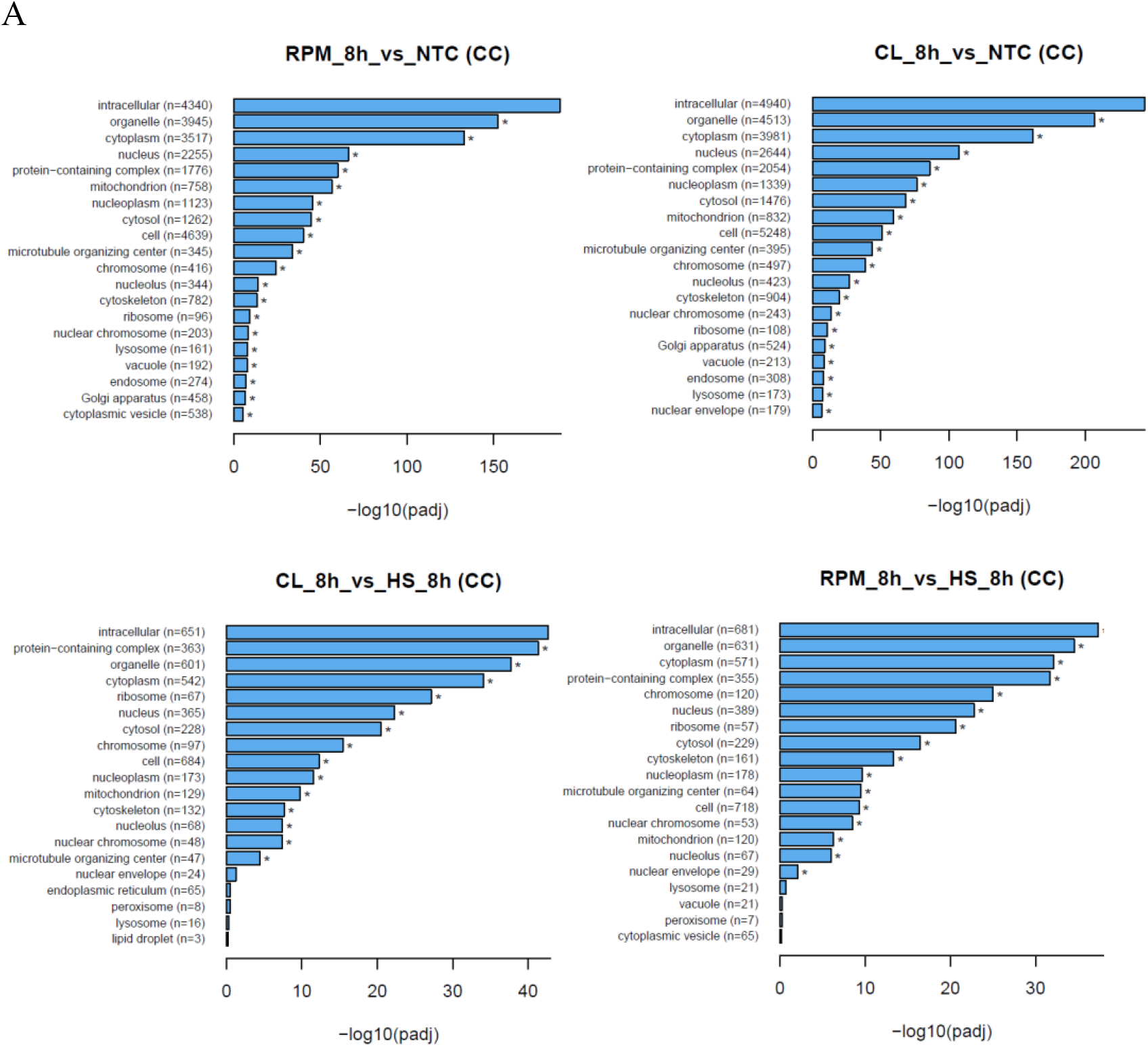
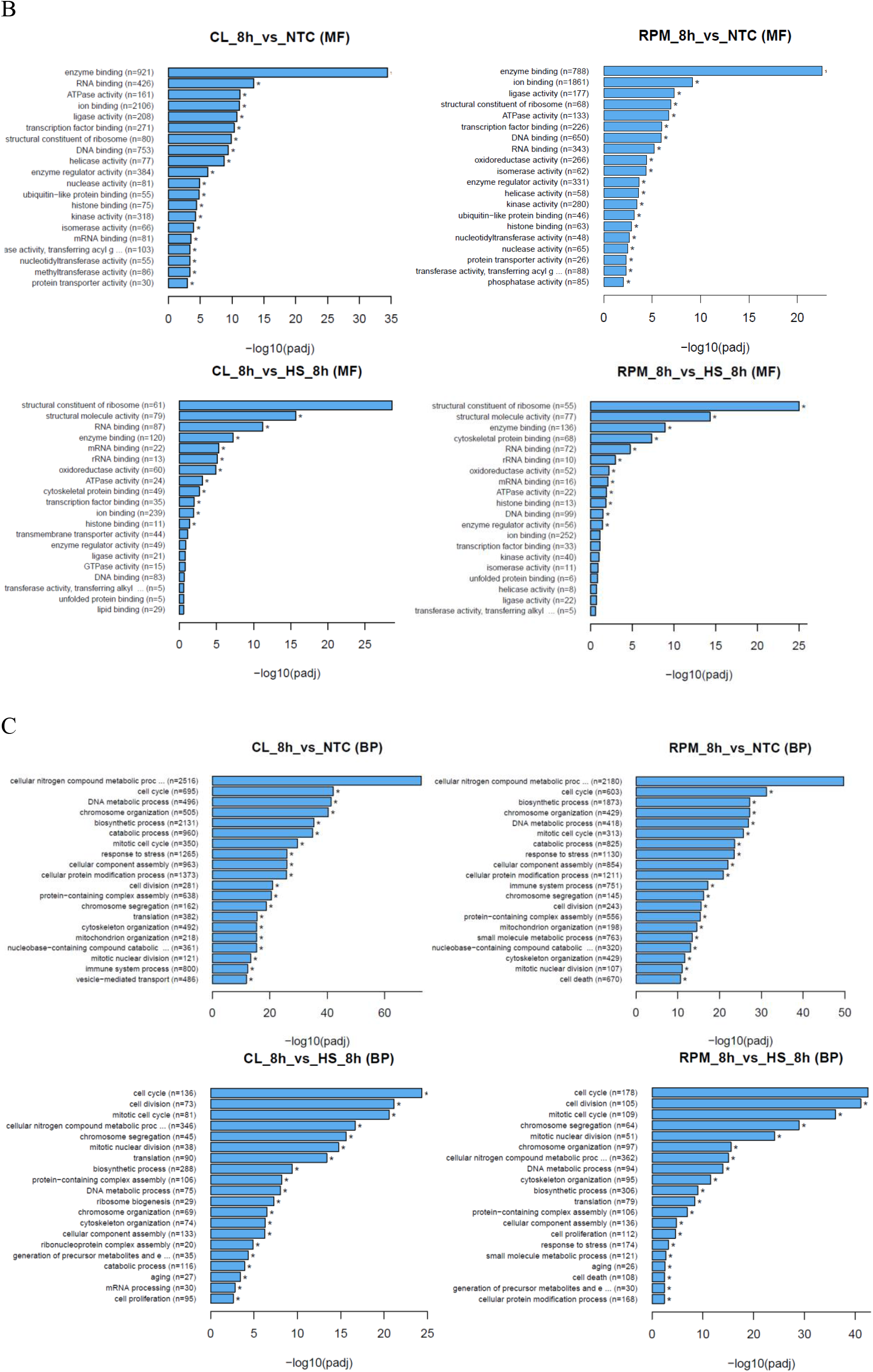
Gene Ontology (GO) enrichment analysis. Treatment versus control comparison for: (A) cellular component (CC), (B) molecular function (MF) and (C) biological processes (BP) terms. Top 20 significantly enriched GO terms are shown.

**Supplementary figure S3.**
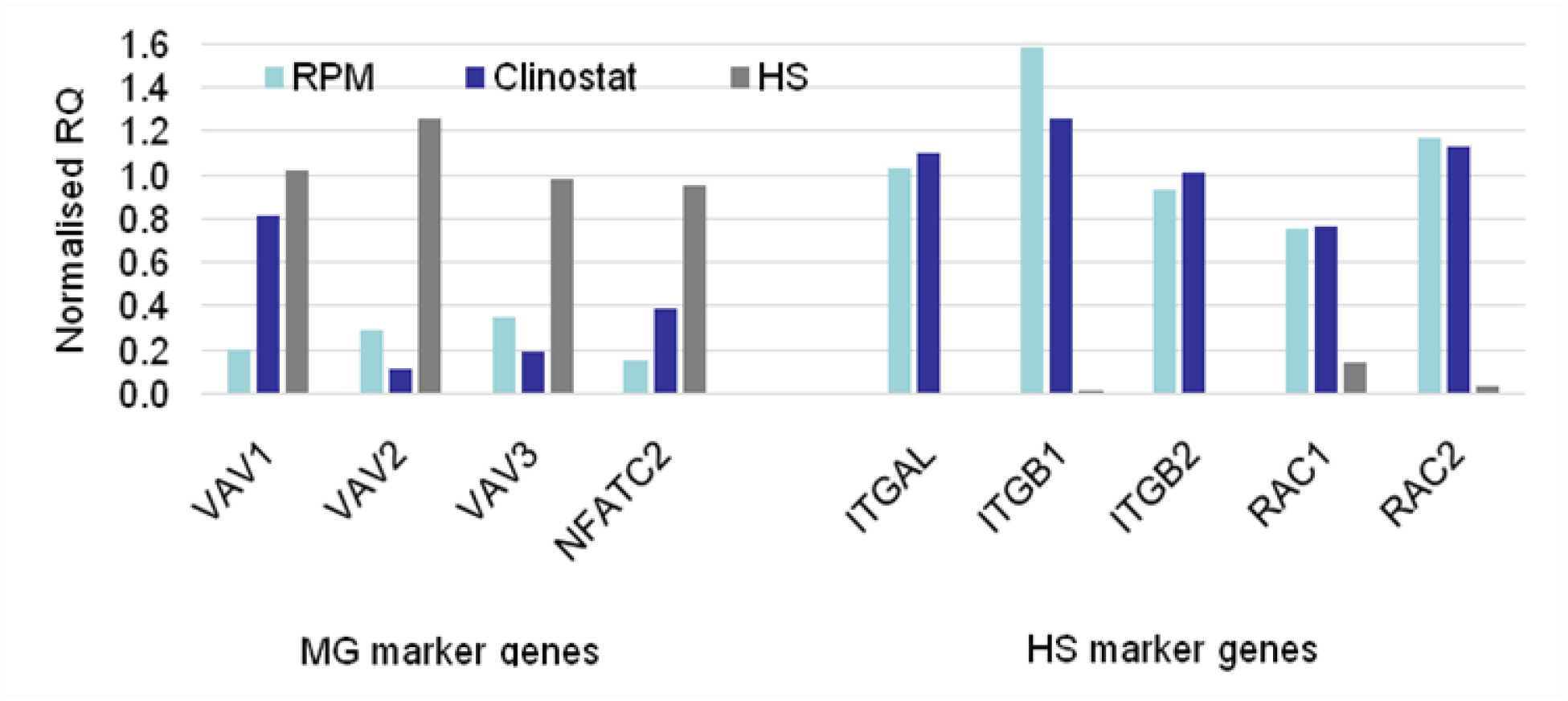
Candidate marker genes discriminating T cell transcriptome responses. Selected differentially expressed genes of actin cytoskeleton network and related genes can be used as potential markers for the response of T cells to simulated microgravity (MG) and to hydrodynamic stress (HS)

## Supplementary Tables

**Table S1.** Three groups of actin cytoskeleton network and related genes according to their relative differential gene expression levels in response of human T cells to simulated microgravity (MG) and under hydrodynamic stress (HS) confirmed by using RT-qPCR.

| Relative gene expression |  |  |
| --- | --- | --- |
| MG > HS | MG = HS | MG < HS |
| CTLA4 | ACTB | ACTG1 |
| EZR | CENPE | FOXP3 |
| ITGAL | MKL1 | MYL9 |
| ITGB1 | MKL2 | NFATC2 |
| ITGB2 | PFN2 | PFN1 |
| RAC1 | SRF | TRP53BP2 |
| RC2 | TLN1 | VAV1 |
| SESN1 | WASF2 | VAV2 |
| TCF12 |  | VAV3 |
| WAS |  |  |

**Table S2.** RNA-seq data Quality Control summary.

| Sample name | Raw reads | Clean reads | Raw_data (G) | Clean_data (G) | Error_rate (%) | Q20 (%) | Q30 (%) | GC_content (%) |
| --- | --- | --- | --- | --- | --- | --- | --- | --- |
| HS_8h_1 | 22275588 | 21763739 | 6.7 | 6.5 | 0.02 | 98.35 | 94.96 | 44.63 |
| HS_8h_2 | 25204052 | 24721513 | 7.6 | 7.4 | 0.02 | 98.22 | 94.63 | 49.22 |
| NTC_1 | 21231374 | 21054307 | 6.4 | 6.3 | 0.03 | 97.79 | 93.58 | 48.93 |
| NTC_2 | 20185658 | 20021323 | 6.1 | 6.0 | 0.02 | 98.21 | 94.53 | 49.63 |
| CL_8h_1 | 24315174 | 23786569 | 7.3 | 7.1 | 0.02 | 98.22 | 94.66 | 49.26 |
| CL_8h_2 | 23844401 | 23296130 | 7.2 | 7.0 | 0.02 | 98.19 | 94.57 | 49.18 |
| RPM8h_1 | 25600529 | 25051563 | 7.7 | 7.5 | 0.02 | 98.20 | 94.61 | 50.06 |
| RPM8h_2 | 22988902 | 22513805 | 6.9 | 6.8 | 0.02 | 98.07 | 94.39 | 48.78 |
| CL_24h | 27021052 | 26760888 | 8.1 | 8.0 | 0.03 | 97.63 | 93.13 | 48.71 |
| RPM24h | 21106398 | 20567883 | 6.3 | 6.2 | 0.02 | 98.22 | 94.61 | 49.56 |

**Table S3.** Overview of RNA-seq data Mapping Status.

Overview of RNA-seq data Mapping Status
| Sample name | Total reads | Total mapped | Multiple mapped | Uniquely mapped | Read 1 | Read 2 | Reads map to '+' | Reads map to '-' |
| --- | --- | --- | --- | --- | --- | --- | --- | --- |
| HS_8h_1 | 43527478 | 41999085<br>(96.49%) | 1920836<br>(4.41%) | 40078249<br>(92.08%) | 20065964<br>(46.10%) | 20012285<br>(45.98%) | 20033591<br>(46.03%) | 20044658<br>(46.05%) |
| HS_8h_2 | 49443026 | 47804795<br>(96.69%) | 1935536<br>(3.91%) | 45869259<br>(92.77%) | 22985377<br>(46.49%) | 22883882<br>(46.28%) | 22924203<br>(46.36%) | 22945056<br>(46.41%) |
| NTC_1 | 42108614 | 40639633<br>(96.51%) | 1341423<br>(3.19%) | 39298210<br>(93.33%) | 19725847<br>(46.85%) | 19572363<br>(46.48%) | 19643692<br>(46.65%) | 19654518<br>(46.68%) |
| NTC_2 | 40042646 | 38799106<br>(96.89%) | 1315043<br>(3.28%) | 37484063<br>(93.61%) | 18781111<br>(46.90%) | 18702952<br>(46.71%) | 18737591<br>(46.79%) | 18746472<br>(46.82%) |
| CL_8h_1 | 47573138 | 46039566<br>(96.78%) | 1665988<br>(3.50%) | 44373578<br>(93.27%) | 22228572<br>(46.73%) | 22145006<br>(46.55%) | 22179441<br>(46.62%) | 22194137<br>(46.65%) |
| CL_8h_2 | 46592260 | 45158442<br>(96.92%) | 1615703<br>(3.47%) | 43542739<br>(93.45%) | 21819540<br>(46.83%) | 21723199<br>(46.62%) | 21763512<br>(46.71%) | 21779227<br>(46.74%) |
| RPM8h_1 | 50103126 | 48551851<br>(96.90%) | 1743876<br>(3.48%) | 46807975<br>(93.42%) | 23452298<br>(46.81%) | 23355677<br>(46.62%) | 23394967<br>(46.69%) | 23413008<br>(46.73%) |
| RPM8h_2 | 45027610 | 43384044<br>(96.35%) | 1601933<br>(3.56%) | 41782111<br>(92.79%) | 20937166<br>(46.50%) | 20844945<br>(46.29%) | 20883276<br>(46.38%) | 20898835<br>(46.41%) |
| CL_24h | 53521776 | 50474422<br>(94.31%) | 1761017<br>(3.29%) | 48713405<br>(91.02%) | 24427089<br>(45.64%) | 24286316<br>(45.38%) | 24348363<br>(45.49%) | 24365042<br>(45.52%) |
| RPM24h | 41135766 | 39828513<br>(96.82%) | 1328387<br>(3.23%) | 38500126<br>(93.59%) | 19287634<br>(46.89%) | 19212492<br>(46.71%) | 19243508<br>(46.78%) | 19256618<br>(46.81%) |

**Table S4.**
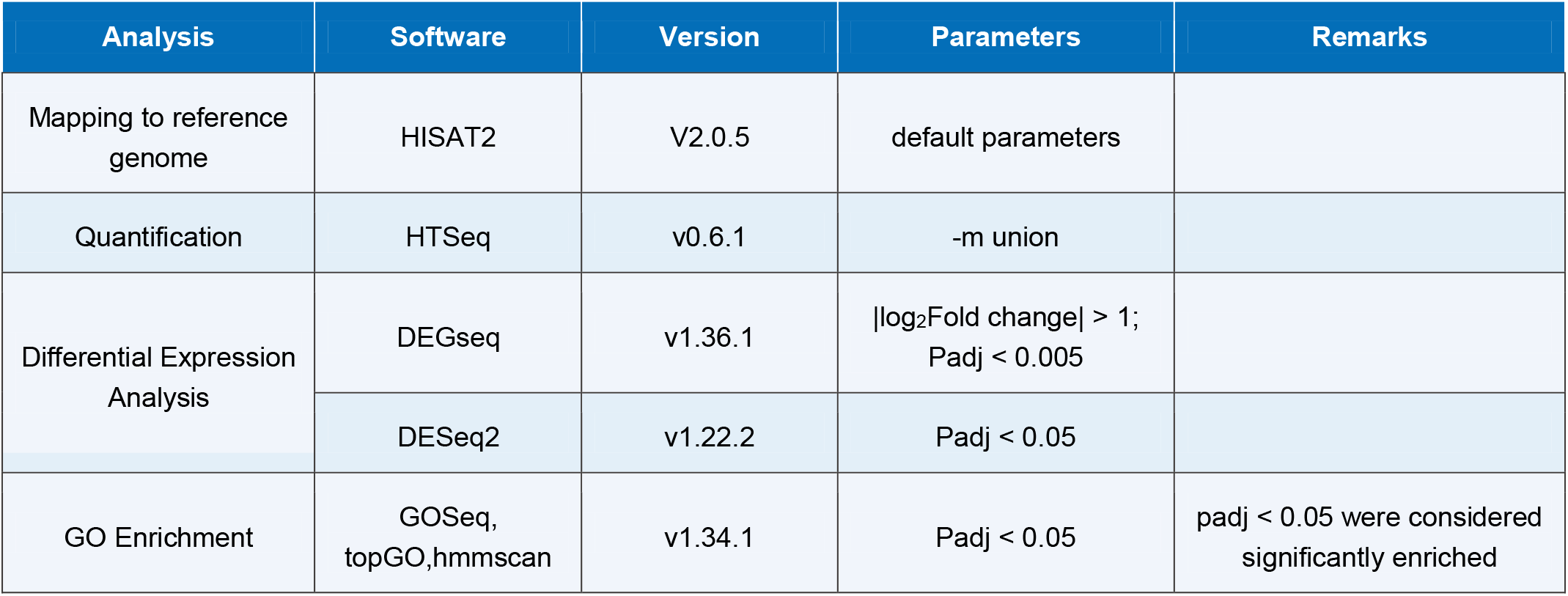
List of software used for RNA-seq data analysis.

